# Toxicity of TiO2, SiO2, ZnO, CuO, Au and Ag engineered nanoparticles on hatching and early nauplii of *Artemia sp.*

**DOI:** 10.1101/329201

**Authors:** R Rohit, Ch. Lakshmi N Murthy, Mohammed M Idris, Shashi Singh

**Affiliations:** CSIR-CCMB (Centre for Cellular and Molecular Biology), Uppal Road, Hyderabad 500 007, INDIA

## Abstract

The potential of environmental release enhances with increased commercial applications of the nanomaterials. A simple and efficient test to estimate the acute toxicity of nanoparticles is carried out in this work using *Artemia* species and their hatching rate. We have tested six different engineered nanoparticles (silver, gold, copper oxide, zinc oxide, titanium dioxide and silicon nanoparticles) and three soluble salts (CuSO4, ZnSO4 and AgNO3) on *Artemia sp*. The physiochemical properties of the nanoparticles involved in this study are estimated and their properties in normal water and marine water were analyzed. Hydrated and bleached *Artemia* cysts were allowed to hatch in continuously aerated, filtered sterile salt water containing nanoparticles; hatching of viable nauplii vs total hatchlings were recorded. In parallel, Standard *Artemia* toxicity test was conducted on the nauplii monitoring the viability. A comparison of results obtained in both experiments is discussed. The toxicity of the nanoparticles was compared and the order of toxicity is estimated as Ag>CuO>ZnO>Au>TiO2>SiO2.

## Introduction

Nanomaterials with their ever expanding diversity, unique properties, and endless applications pose risk to environment and human health. There is dearth of information on the impact or the risk of nanoscale objects to the environment. Increasing utility would mean enhanced exposure of the ecosystems to these unknown risks. The release into environment may start with production, during applications, by weathering and finally wastes.

Nanomaterials can be designed from almost any material – metal/oxides, carbon, organic, biomaterial and in any form, by strictly adhering to the size range of nanometer at least in one dimension. Nanomaterials behave differently from the bulk due to their unique physico-chemical properties. Information on predictability of their behaviour or state in due course is almost negligible. Not only there is limited information on bioavailability and biopersistence of these particles, the behaviour of the particles in various conditions pH, salinity and other biotic factors are not clearly known [1, 2]. Still efforts are being made to identify the risks associated with the nano-material applications.

Hazard identification is an important step for risk assessment of nanomaterials. Most of the methodologies used in hazard identification are the ones used for chemicals in general. Being different from bulk material, the nanoforms of the same materials may pose some restraints or advantages over the use of methodologies.

Aquatic systems are the inevitable receptacles for the materials released in the environment. The organisms therein get exposed to the nanomaterials accumulated from seepage and flow through. Many assays for ecotoxicology have been tested for nanomaterials - in bacteria [3], fishes and fish embryos [4, 5]; copepods [6]; Daphnia [7,8]. In this study *Artemia sp*. are used to study the nanomaterial toxicity. Artemia is a non-selective filter feeder organism which available worldwide in highly saline waters. Due to their ubiquitous distribution, robustic nature and cost effective culture they make a good model organism for toxicological assays. Though artemia are considered to be one of the insensitive models for a lot of chemicals the early developmental stages of Artemia are highly vulnerable to many of the test materials. Few studies are reported for nanomaterial toxicity using *Artemia sp*. Also [9-13]. Most of the studies are carried out in newly hatched nauplii that are exposed to varying concentration of nanomaterial; Hatching of hydrated cysts has not been used as an end point for toxicity except in [14] though some earlier studies had shown sensitivity of hatching rate of Artemia to metals [15-17].

In this paper we have studied acute toxicity of 4 metal oxide nanoparticles (TiO2, SiO2 and CuO) and two metal nanoparticles (Ag and Au) to nauplii of Artemia and on the hatching of their hydrated cysts. Mortality and hatching rates are considered as end points respectively for the naupili test and hatching. Both the tests show good inverse correlation in sensitivity to nanoparticles, thereby implying another bioassay can be developed for hazard assessment of nanoparticles.

## Materials and Methods

### Preparation of nanoparticles and characterization

All the nanoparticles used in the study are obtained from the consortium NANOVALID project. Silicon nanoparticles were supplied by Nanologica AB, Silver nanoparticles were obtained from Colorobbia; Gold nanoparticles from INMETRO., Copper oxide nanoparticles from Intrinsiq Materials (UK)., Zinc oxide nanoparticles from Nanogate (Germany) and Titanium dioxide particles were synthesized in house using the existing protocol [18]. A suspension of eNPs is made by suspending sio2, Tio2, Cuo and Zno NPs at a concentration of 2mg/ml. The suspension is sonicated at ∼ 30W with pulse of 50% for 10 minutes. Silver nanoparticles are obtained as suspension at a concentration of 4% with PVP as a stabilizing agent, gold nanoparticles in water at a concentration of 0.006%. Working dilutions of the nanoparticles are made in salt water (NaCl) for various studies.

All of these nanoparticles used in this study are characterized in the NANOVALID consortium in round robin manner. TEM (transmission electron microscopy) and DLS (dynamic light scattering) are used in this study for basic characterization of nanoparticles before every experiment. In TEM studies the nanoparticle suspension of 50µg/ml was applied to formvar coated copper grids and air dried. The grids were examined in JEOL 2010 TEM at 100KV using 20µ aperture, about 10 images were taken using GATAN camera and images were analysed using GATAN software. The sizes of particles were used for analysis. The nanoparticle suspensions at concentration of 25µg/ml in milliQ water and salt water were measured in Dynamic light scattering system (Horiba Nanopartica SZ100). Each suspension was analyzed thrice for three days and the data was represented as mean ± SEM. Solvents used to prepare the nanoparticles were measured for size determination as controls.

### *Artemia* cultures

For every 1 litre culture, 1.0 gram of *Artemia* cysts were used (San Francisco Bay strain). These cysts are hydrated in distilled water under aeration for 45 minutes and then bleached to decapsulate the cysts wall using 20% sodium hypochlorite solution for 15 minutes. The bleached *Artemia* eggs were then washed for 6 times with distilled water and allowed to hatch in 25% salt water (pH 8.0) with continuous light and aeration for 24 hours [19]. The temperature was maintained at 28°C which is the optimum temperature for hatching of *Artemia* eggs. Nanoparticles are added to the culture in concentrations of 100mg/l and 10mg/l.

### Toxicity on hatching rate

Five types of nanoparticles (Ag, Au, SiO2, TiO2, CuO and ZnO) are tested for their toxicity on brine shrimp cultures. The *Artemia* eggs are allowed to hatch under standard culture conditions and are exposed to two different concentrations of each nanomaterial (100mg/L & 10mg/L). The hatching rate, mortality and viability are checked after 24 hours for each

*Artemia* cultures exposed to nanoparticle. Three aliquots of 100µl was taken from each flask after thorough mixing and counted under the stereo-microscope. Number of hatched *Artemia* (immotile vs. motile), umbrella stage and unhatched cyst were counted. Hatching rate was expressed as percent of total number of fully hatched *Artemia* in comparison to total *Artemia* in the aliquot. The experiment was repeated six times. The average and SEM was calculated for each sample.

### Standard Artemia toxicity Test

In a second set of experiments, following protocol of *Artemia* hatched under normal conditions in salt water [20] were exposed to nanoparticles. 10 nauplii were transferred to six well plates and the nanoparticles at 100mg/l, 10mg/l, 1mg/l, 0.1mg/l and 0.01mg/l concentrations were added. The viability of the larvae was counted at intervals of 6, 12, 18, 24, 30 and 48 hours. Mortality of the larvae was used as end point. Mortality was recorded as cessation of swimming or any movement by nauplii. LD50 was calculated for each particles using Graphpad software (www.graphpad.com/quickcalcs/).

### Oxidative stress test

Stresses due to reactive oxidation species can be observed by treating the naupili exposed to different concentrations with10µM DCFDA (dichloroflouresciendiacetate). DCFDA which is a chemiluminiscent probe reacts with ROS generated due to oxidative stress and the fluorescence can be imaged using microscope. The nanoparticle treated naupili exposed to DCFDA are imaged in Zeiss Axiovert imager.

### Validity of results

The hatching studies have been conducted 6 times and the validity criterion used is, low mortality in the control. The data was analyzed for statistic validation using Graphpad software (www.graphpad.com/quickcalcs/). In the *Artemia* toxicity tests the validity criterion is mortality below 10% in 24 hr period. The correlation test was performed for both sets of data using on line software for correlation coefficient and comparing data in Microsoft excel.

## Results

### Analysis of nanoparticles

The size of nanoparticles both in distilled water and salt water were measured. The nanoparticles showed reduced size in high salt concentration by Transmission Electron Microscopy. The transmission electron microscope studies show no apparent change in aggregation or dispersion of particles (Figure 1). Nanoparticles in water showed a higher DLS value for all the nanoparticles. Nanoparticles exhibited different behaviour in salt water; Silver, TiO2, CuO and ZnO nanoparticles showed higher values. Gold and silicon dioxide had DLS values lesser as compared to particles in Milli Q water. (Figure 2). Visible flocculation was observed in case of Copper and Zinc oxide nanoparticles. DLS measurements for these nanoparticles gave very high values corroborating the flocculent seen (Figure 2).

**Figure 1:**
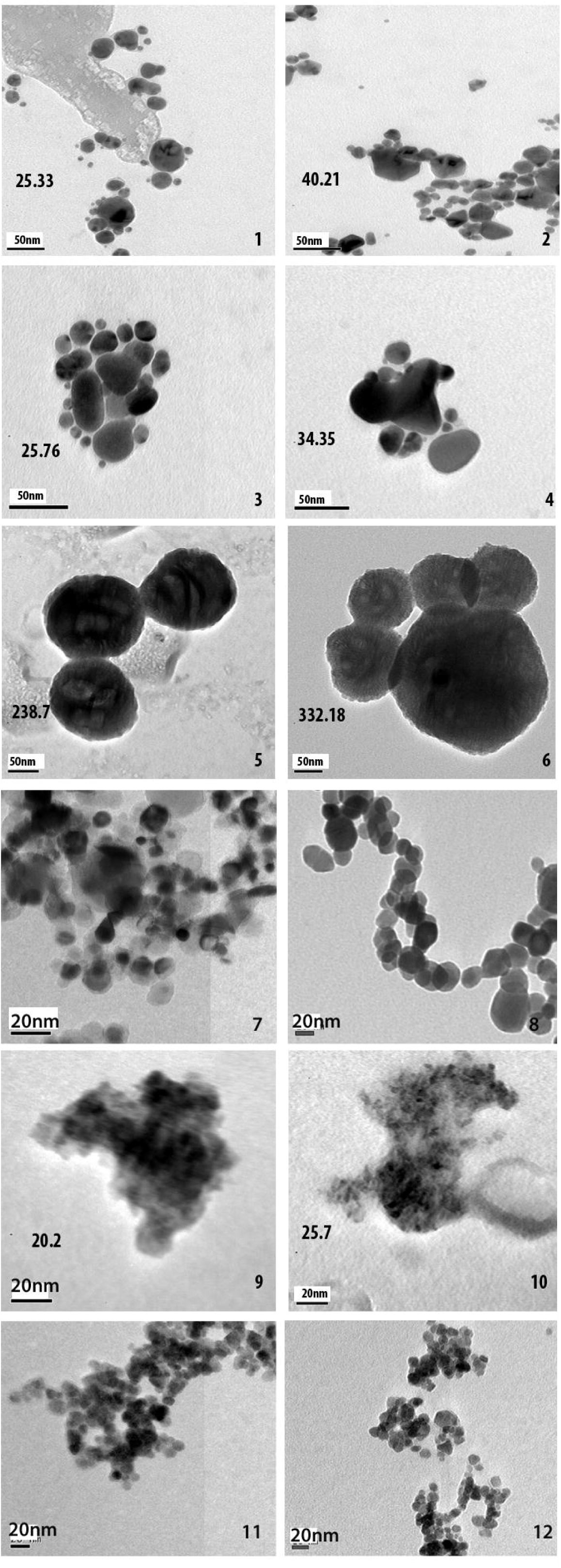
Transmission Electron micrograph of nanoparticles suspended in distilled water and high salt solution. Panel on the left represents particles in salt water and on right particles in DW. The particles are silver eNP (1, 2); gold (3, 4); Silicon (5, 6) CuO (7, 8) Titanium dioxide (9, 10) and ZnO nanoparticles (11, 12).

**Figure 2:**
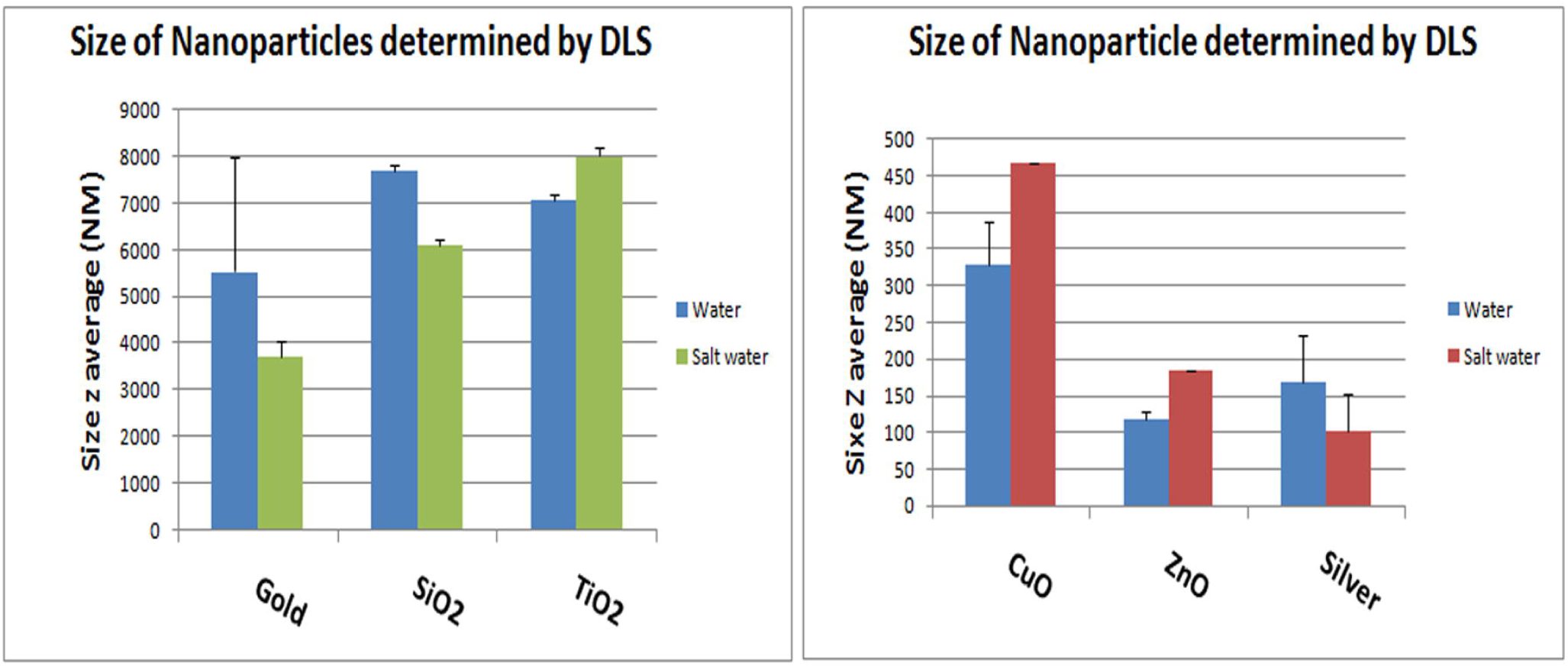
Size distributions of nanoparticles in distilled water and salt water. Many nanoparticles show reduced particle size due to dispersion in salt water. Copper Oxide and Zinc Oxide that show higher values also show visible flocculation when suspended in salt water.

The nanoparticles suspended both in Milli Q water and 25%NaCl solution (salt water) were measured up to 48 hours to see change in their dispersion pattern (Figure S1). Silver NP, Titania and silica did not show much difference in dispersion in Milli Q water over three days but on third day the particles showed higher aggregate sizes in salt water. In CuO and ZnO NPs visible flocculation and sedimentation was seen in case of both Milli Q and salt solution by third day.

### Exposure studies

Brine shrimp were allowed to hatch in filtered sterile salt water kept at continuous aeration that also keeps the particles in suspension throughout the experiment. Brine shrimp normally showed a hatching rate of 74%± 5.5, the viability of these hatched nauplii was about 97%. The hatching rate of brine shrimp altered drastically in presence of nanoparticles (Table 1; Figure 3). The hatching rate dropped with increase in concentration of nanoparticles in salt water. The highest effect was seen in presence of silver and copper oxide nanoparticles. The hatching rate dropped to about 29% in the two nanoparticles; of these only 50% were motile and viable at a concentration of 100mg/L (Table 1). The hatching rate remained lower at 10 mg/L of silver nanoparticles but the viability of hatched nauplii was good at 85%. In copper oxide also hatching rate was low with better viability around 64%. Similarly, with silver nitrate or in presence of capping agent PVP the hatching slightly improved (33% and 41% resp.) with good viability of 70 and 84%, and about 60% in case of copper sulphate with viability of 33%.

**Table 1:** Hatching rate of Artemia cysts and immediate mortality of the nauplii.

| Hatching | Percent Hatching rate | SEM | p value | Immediate Mortality of hatchlings | SEM | p value |
| --- | --- | --- | --- | --- | --- | --- |
| Blank | 74.6 | 5.176899 |  | 2.026 | 1.3 |  |
| Agnp100 | 29.3 | 9.975868 | 0.013 | 48.3075 | 14.51 | 0.0105 |
| Agnp10 | 32.2 | 8.239192 | 0.010 | 15.3525 | 5.0 | 0.0329 |
| AgNO3 | 33.4 | 9.19874 | 0.019 | 30.7125 | 12.1 | 0.0426 |
| PVP | 41.1 | 9.95419 | 0.079 | 15.705 | 5.621 | 0.0462 |
| au1 | 57.1 | 10.40615 | 0.622 | 4.3125 | 1.833 | 0.46 |
| 0.5au | 53.1 | 9.393431 | 0.378 | 9.0825 | 2.8 | 0.0698 |
| SiO<sub>2</sub>-100 | 48.9 | 5.603092 | 0.390 | 20.037 | 11.9 | 0.1761 |
| SiO<sub>2</sub>-10 | 63.8 | 9.656019 | 0.943 | 8.177 | 5.8 | 0.3646 |
| TiO<sub>2</sub>- 100 | 41.51 | 12.67946 | 0.147 | 32.5 | 11.4 | 0.0277 |
| TiO<sub>2</sub>- 10 | 54.3 | 5.480797 | 0.275 | 18.41 | 4.8 | 0.1587 |
| TiO<sub>2</sub>- 1 | 59.07 | 6.012516 | 0.631 | 2.2 | 0.7 | 0.0616 |
| CuO-100 | 29.14253 | 13.25419 | < 0.0001 | 55 | 39.02456 | 0.0077 |
| CuO- 10 | 43.23553 | 17.32773 | < 0.0001 | 33.816 | 7.411374 | 0.0536 |
| CuSO<sub>4</sub> | 59.05675 | 11.07297 | 0.0078 | 47 | 21.74665 | < 0.0001 |
| ZnO-100 | 60.52844 | 13.36209 | < 0.0001 | 49.6 | 32.5 | 0.0018 |
| ZnO-10 | 63.38297 | 14.96577 | < 0.0001 | 33.75 | 30.13857 | 0.0119 |
| ZnSO<sub>4</sub> | 44.71964 | 22.27028 | < 0.0002 | 42 | 37.96929 | 0.0139 |

**Figure 3:**
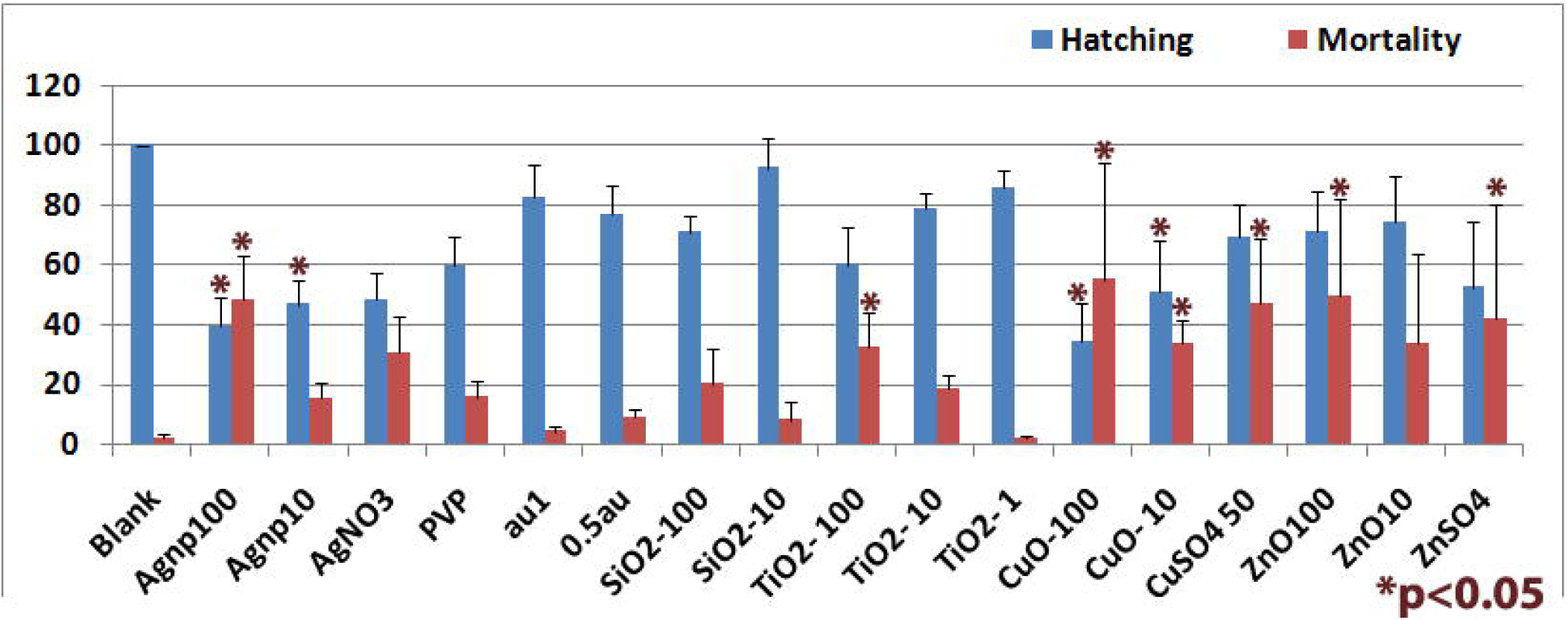
Hatching rate and percent of dead nauplii in the hatching experiment. Silver nanoparticles show the highest toxicity both in terms of hatching and immediate survival of nauplii after hatching. Other nanoparticles show decline in hatching but posthatching survival is not affected. Silicon nanoparticles at lower concentration do not affect hatching and survival. * significance P<0.05.

Hatching rate was low in presence of other nanoparticles (gold, silicon, titanium, copper and zinc oxide) at around 50-60% as compared to control (Table 1; Figure 4) and post-hatching mortality was not very high. In presence of zinc oxide nanoparticles, the hatching rate was high at 60% but the cysts had a tendency to clump together. In titanium dioxide at 100mg/L, hatching rate was 41.50% with 67.8 % viability; hatching rate was better in 10mg/L at 54% showing ∼90% viability. Gold and silicon dioxide nanoparticles displayed very good viability after hatching (Figure 3). SiO_2_ nanoparticles at a concentration of 10mg/L displayed hatching rate and viability equivalent to the controls.

**Figure 4:**
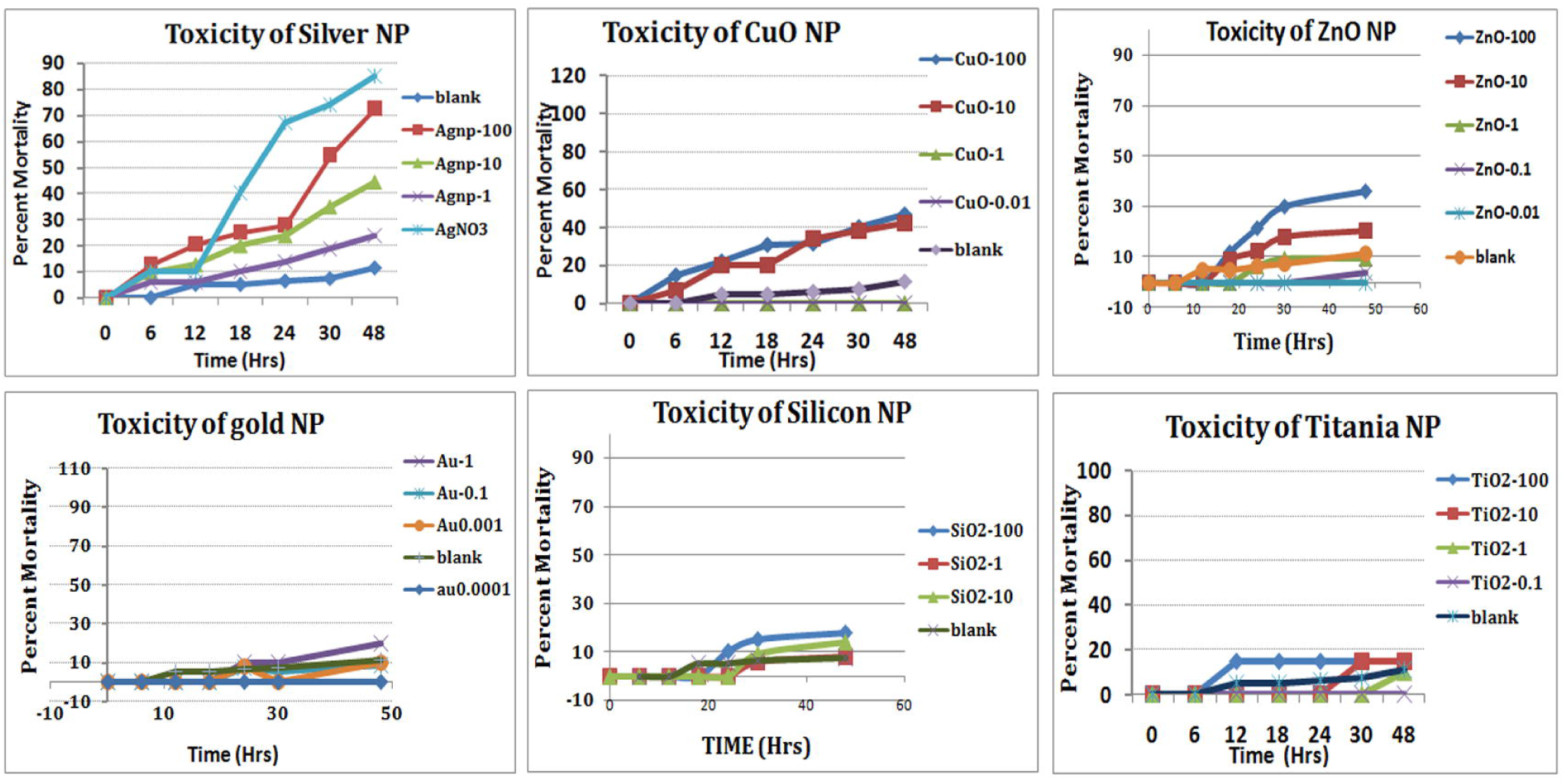
Mortality of nauplii in the standard ARC test. Silver nanoparticles show the highest toxicity in concentration dependent manner with LC50 of 10.12mg/L at 24 hours, silver in ionic form and the supernatant of nanoparticles show high toxicity. Other nanoparticles show concentration dependent mortality, with LC_50_ of CuO, Silica and Titania was around 20mg/L.

The pH of salt water did not change upon addition of nanoparticles and remained around 6.3-up to 48 hours. Hatched nauplii appeared normal and swimming vigorously. Many of the nauplii had emerged but were immotile/dead; some were still-dead and trapped in the membrane (Figures S2.1-2.6). There were no changes in morphology of nauplii hatched in presence of nanoparticles. Silver and copper oxide nanoparticles with maximum toxic effect on hatching rate also did not show any gross morphological abnormalities. Staining of hatched *Artemia* with DCFDA for signs of stress showed staining for ROS in few still trapped nauplii and the gut of dead nauplii (Figure S3).

### Larval toxicity test

Following the ARC (*Artemia* Reference Centre) test protocol, the larvae hatched in salt water were exposed to nanoparticles for up to 48 hours and the viability recorded every 6 hours for first 24hours. *Artemia* were not fed during the test period. The nauplii showed variability in viability to various nanoparticles. The most toxic was the silver nanoparticles and silver in ionic form (Table 2). Silver in ionic form was highly toxic as 80% of the nauplii were dead in 24 hours in silver nitrate solution (Figure 4). The LC_50_ of the silver nanoparticles was around 10.12mg/L at 24hours. Other nanoparticles used in the study were nontoxic initially but after 30 hours (Table 2), Many *Artemia* perished in concentration dependent manner (Figure 4). LC_50_ of gold was 2.53E-03mg/Lat 48h. LC_50_ of CuO, Silica and Titania was 20.92; 23.59 and 18.94mg/L at 48 hours. In presence of zinc oxide nanoparticles, around 36% nauplii perished in 48 hrs in the highest concentration used with the LC50 above 100mg/L.

**Table 2:** Lethal concentration of nanoparticles for larval toxicity test.

| Nanoparticle(concentration) | 24hour LC50% | 48h hour LC50% |
| --- | --- | --- |
| Ag | 10.12mg/l |  |
| Au |  | 2.5*10<sup>-3</sup>mg/l |
| SiO<sub>2</sub> |  | 23.59mg/l |
| TiO<sub>2</sub> |  | 18.94mg/l |
| CuO |  | 20.92mg/l |
| ZnO |  | 100mg/l |

In the ARC test; we do see sedimentation and flocculation of nanoparticles by 48 hrs. The nauplii maintained in salt water containing nanoparticles showed accumulation of eNPs in the gut of *Artemia* by end of 24 hours (Figure 5). In case of silver nanoparticles, the nauplii that are dead within 24 hours, the gut does not show accumulation of material (Figure 5), even the nauplii appear stunted. Staining of the nauplii exposed to nanoparticles show signs of stress when stained with DCFDA for ROS and DHE for Superoxide (Figure 5). Nauplii exposed to silver nanoparticles show highest staining, followed by titania and silicon. Nauplii exposed to other nanoparticles do not show any signs of ROS or Superoxide but had accumulation of nanoparticles in the gut (Figure 5).

**Figure 5:**
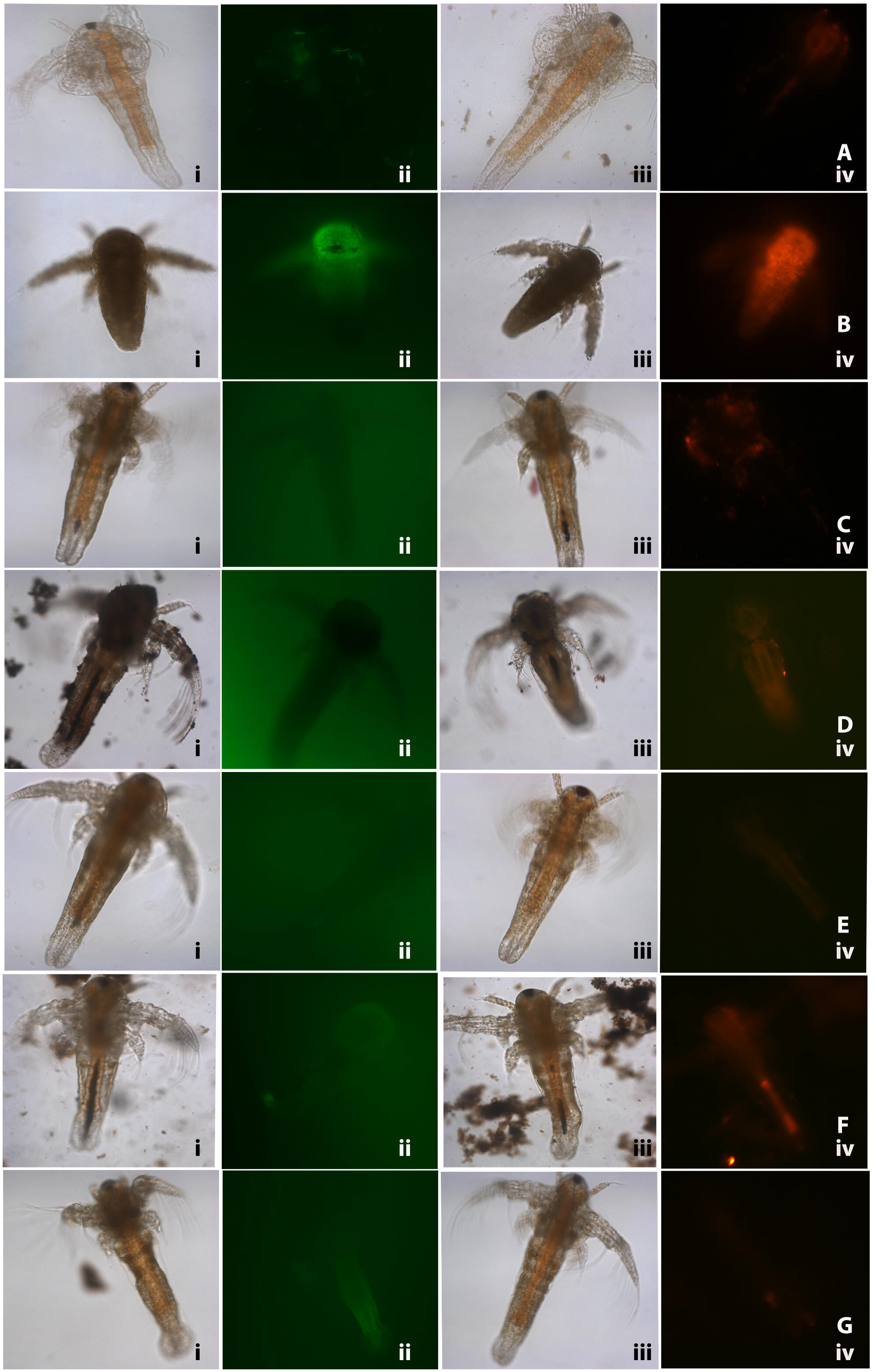
Nauplii of *Artemia* in standard ARC test stained with DCFDA (green) for ROS and DHE (Red) for superoxide accumulation due to stress. Nauplii exposed to silver NPs, TiO_2_ and silicon NP show higher staining for both ROS and superoxide at 24 h. Row A-control, B-Silver NP, C-Gold NP; D-CuO NP; E-ZnO –NP; F-Titania and G-silicon NP. Some Particles show accumulation in gut.

## Discussion

In this study, we have compared hatching rate of *Artemia* cyst in presence of nanoparticles and the toxicity of nauplii using standard *Artemia* micro-well test at various concentrations of nanoparticle suspension in high salinity conditions. We found a strong correlation in hatching rates and larval mortality in various nanoparticle suspensions (Figure 6).

**Figure 6:**
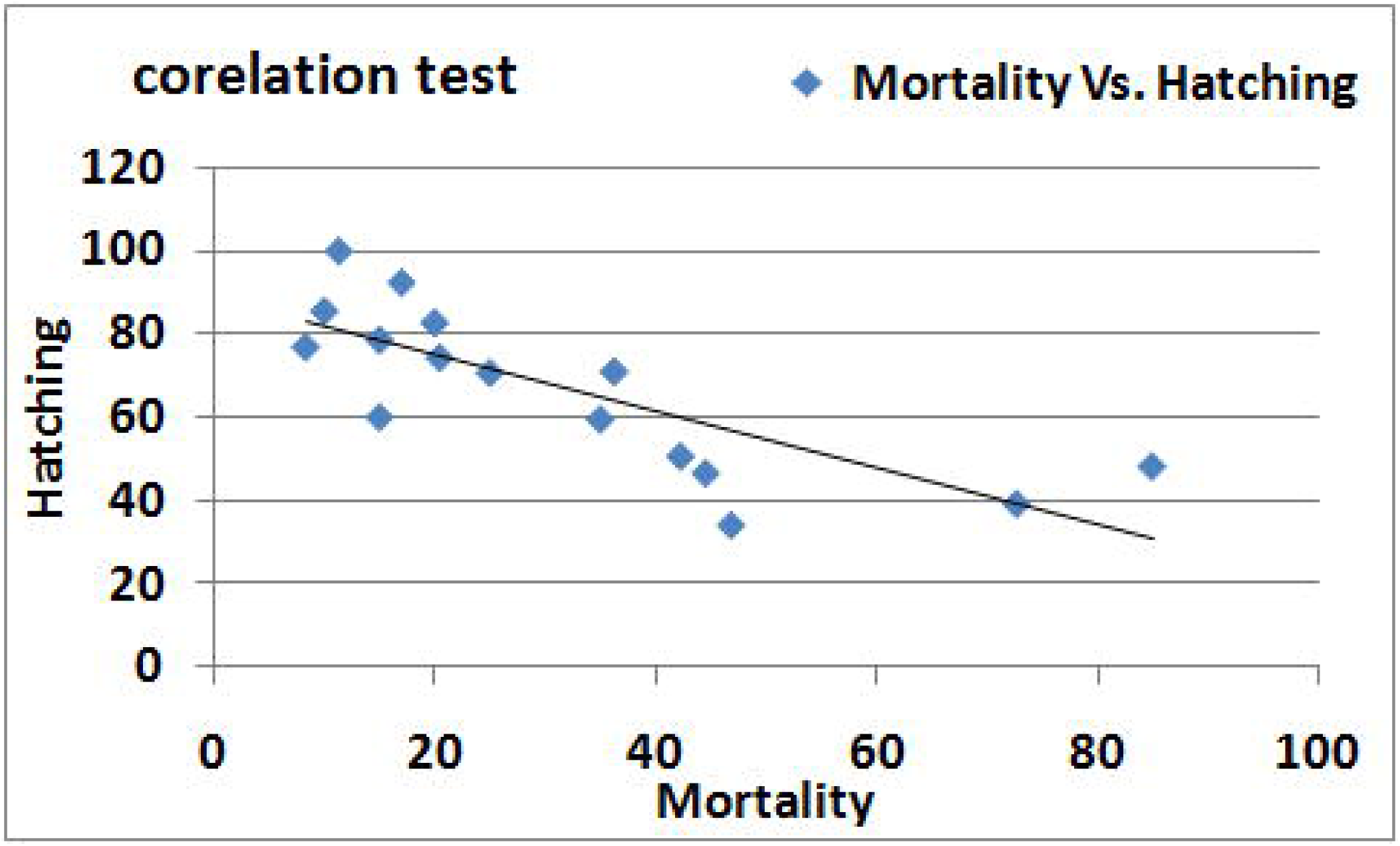
Regression analyses to show correlation between the two sets of results. There is a strong inverse correlation between hatching rate and the mortality in the standard Artemia test (coefficient of correlation −0.748).

*Artemia* are able to tolerate high salinity and adapt to extreme conditions. They have wide distribution worldwide and the cysts can be easily collected. *Artemia* cysts are dormant gastrula states during development encased in hard outer shell; formed under unfavourable conditions. Under simple laboratory conditions, the cysts can be hatched in high salinity water when by about 20 hours the umbrella stage emerges and yields free swimming Nauplii [21, 22]. In the hatching test, we counted the number of umbrella stage and nauplii after 24 hours, among the hatched larvae, number of motile nauplii were recorded for each tested nanoparticle. Hatching or emergence of the nauplii has been used as a sensitive bioassay to study the molecular and physiological effects of metals [15, 23]. Lower hatching rate equates with the toxicity of test compound.

Silver and copper oxide nanoparticles show high toxicity among the various particles tested in concentration dependent manner. Their toxicity could have been due to ionic forms so we have centrifuged the nanoparticle suspension and resuspended in brine solution. These resuspended particles can be seen as nanoparticles even after 72hours in high salinity water and cause toxicity by affecting hatching rates and viability both. Though ionic silver precipitates out due to high salinity, it still affected the hatching and subsequent viability of the larvae. Silver nitrate solution and copper sulfate representing ionic forms also showed high toxicity. Other nanoparticles like; gold, titanium dioxide and SiO2 nanoparticles had some effect on hatching rates and survival after hatching in concentration dependent manner. Silicon nanoparticles at low concentration were very conducive in fact showed better hatching as compared to control. Hatching rates were not much different at 20 hours and 24 hours but the mortality of hatched nauplii was higher in some nanoparticles at 24 hours (data not shown). Locomotion of the hatched nauplii was also affected in presence of nanoparticles.

In the standard *Artemia* test, nauplii hatched in salt water are exposed to nanoparticle suspensions. The mortality and toxic effects expressed as LC_50_; the concentration of an agent at which 50 per cent of the tested animals are dead after 24 hr; were chosen as criteria of the toxicity [24]. The standard *Artemia* toxicity test results were comparable and silver nanoparticles ranked very high in toxicity, followed by copper and zinc oxide nanoparticles. Titanium dioxide, gold and silicon nanoparticles in that order, showed a better survival of nauplii up to 48 hrs. On comparing the two methods, we found a very good inverse correlation between the results obtained on effects of NPs on hatching rates and mortality test on the hatched larvae. The particles that reduced the hatching rate showed an increase in mortality in the ARC test using the normally hatched nauplii. The correlation test showed a coefficient of −0.7853. Our results correlate with some other toxicity assays performed on *Artemia* using nanoparticles [25, 26, 11]. Comparing the viability of nauplii (inverse of mortality) in ARC test with the hatching rate showed no statistically significant difference in the mean values for each concentration of different particles (P=0.683) in the paired T test.

The advantages of these assays are that they are easy to perform, short term and use mortality as end point for assessing toxicity of the test material. The question of altered solubility may not arise here but sedimentation of these suspensions may vary for different particles. So far we have seen that nanoparticles tested in this work remain stable in seawater up to one week, though the ionic or dissolved component may settle down due to precipitation. The added advantage of continuous aeration in the hatching test keeps the particles in suspended form.

Organisms in aquatic environment like daphnia incorporate eNPs via gut. Artemia is also a filter feeder and the NPs may enter the guts of *Artemia* through ingestion. The mortality observed may be due to the uptake of the eNPs clogging the gut as most of the nauplii in presence of eNPS show presence of particles in the digestive tract. In the hatching test, the dead *Artemia* and partially hatched cysts show increased stress in presence of nanoparticles as indicated by DCDFA staining. Even the nauplii exposed to some nanoparticles do show accumulation of ROS and superoxide especially with silver nanoparticles. But with remaining nanoparticles it must be the NP clogging in the gut that could cause mortality.

*Artemia* with capacity to homeostasis in variable external salt concentrations certainly appears to be a model organism to assess the nanoparticle behaviour and ecotoxicity. Suitability of *Artemia* for toxicity tests was established and is often used in pharmaceutical industry and for metal and radiation toxicity [27-29]. *Artemia* hatchability test along with standard lethality/toxicity test definitely could also be used as pre-screening test for nanoparticle toxicity prior to validating it for in vivo toxicity studies. These acute toxicity tests eliminate the need of laboratory maintenance of test species that take up space and time. Use of cysts which are available in enormous quantities as batch also cut down on the variability. With strictly laid out protocol and test conditions, this assay can be adopted for the hazard identification of nanomaterial and high through put methods.

## Acknowledgments

This work was supported by FP7-NANOVALID (Grant No. 263147).

